# miRNA and mRNA Signatures in Human Acute Kidney Injury Tissue

**DOI:** 10.1101/2023.09.11.557054

**Authors:** Danielle Janosevic, Thomas De Luca, Ricardo Melo Ferreira, Debora L. Gisch, Takashi Hato, Jinghui Luo, Yingbao Yang, Jeffrey B. Hodgin, Pierre C. Dagher, Michael T. Eadon

## Abstract

Acute kidney injury (AKI) is an important contributor to the development of chronic kidney disease (CKD). There is a need to understand molecular mediators that drive either recovery or progression to CKD. In particular, the role of miRNA and its regulatory role in AKI is poorly understood. We performed miRNA and mRNA sequencing on biobanked human kidney tissues obtained in the routine clinical care of patients with the diagnoses of AKI and minimal change disease (MCD), in addition to nephrectomized (Ref) tissue from individuals without known kidney disease. Transcriptomic analysis of mRNA revealed that Ref tissues exhibited a similar injury signature to AKI, not identified in MCD samples. The transcriptomic signature of human AKI was enriched with genes in pathways involved in cell adhesion and epithelial-to-mesenchymal transition (e.g., *CDH6, ITGB6, CDKN1A*). miRNA DE analysis revealed upregulation of miRNA associated with immune cell recruitment and inflammation (e.g., miR-146a, miR-155, miR-142, miR-122). These miRNA (i.e., miR-122, miR-146) are also associated with downregulation of mRNA such as *DDR2* and *IGFBP6*, respectively. These findings suggest integrated interactions between miRNAs and target mRNAs in AKI-related processes such as inflammation, immune cell activation and epithelial-to-mesenchymal transition. These data contribute several novel findings when describing the epigenetic regulation of AKI by miRNA, and also underscores the importance of utilizing an appropriate reference control tissue to understand canonical pathway alterations in AKI.

## Introduction

Acute kidney injury (AKI) is an important contributor to chronic kidney disease (CKD) development and mortality.^1-4^ AKI in the clinical setting is heterogenous and available diagnostic tests (e.g., serum creatinine, urinary biomarkers) lack the specificity to delineate AKI etiologies and are not reflective of underlying pathobiology.^5^ Overall, limiting supportive care as sole treatment for AKI, and have impeded improvements in overall morbidity and mortality. Understanding the epigenetic regulation of mRNA by miRNA in renal injury and recovery processes at the target mRNA level, and its transcriptomic regulation may facilitate biomarker discovery to enable early treatment and improve renal survival.

AKI is a multiphasic syndrome, and the molecular characterization of its phases may hold important therapeutic implications.^6; 7^ Using a multiomic approach (i.e., bulk kidney and single-cell RNAseq, proteomics) with a murine model of sepsis-associated AKI (sAKI), we identified a multiphasic renal injury response involving sequential activation of inflammatory and antiviral pathways leading to a global shutdown of protein translation and consequent organ failure, then subsequent recovery.^8; 9^ Recently, the merged Kidney Precision Medicine Project (KPMP) and human biomolecular atlas project (HuBMAP) and single-cell RNAseq (scRNAseq) kidney atlas revealed an adaptive cell state as an important mediator of CKD progression in individuals with AKI and CKD.^10^ This cell state is representative of the equilibrium between successful and failed repair, the latter of which leads to fibrosis.^11-14^ Although the epigenetic regulation of the adaptive cell state by DNA methylation or histone modification has been explored;^15^ regulatory involvement of non-coding RNAs such as microRNAs (miRNA) have yet to be integrated into a comprehensive molecular understanding of this physiological response.

miRNAs regulate cellular transcriptional processes by complementary base pairing to the 3’-untranslated region of target mRNAs, largely promoting rapid degradation of RNA and inhibition of protein translation.^16^ miRNAs are known to regulate pathways of inflammation, immunity, replication and metabolism in AKI.^17-20^ Further study of miRNA-mRNA regulatory networks in AKI will likely broaden our understanding of AKI pathobiology, but also serves to elucidate additional biomarkers of disease activity. miRNAs are readily found and resistant to degradation in body fluids and are reflective of underlying renal organ functions,^21-26^ and thus, have the potential to be valuable molecular diagnostics and therapeutic targets in human AKI.

In this study we sought to describe the miRNA-mRNA regulatory networks in human kidney biopsy samples that may provide insight into adaptive and maladaptive repair processes in AKI. Herein, we performed miRNAseq and mRNAseq on human biobanked biopsy specimens from individuals with the primary histopathological diagnosis of AKI from either acute tubular necrosis (ATN) or acute interstitial nephritis (AIN). The expression signatures of the AKI renal biopsy specimens were compared to reference nephrectomies obtained from deceased kidney donors and unaffected portions of tumor nephrectomies. Because nephrectomy tissues (Ref) are often affected by some form of hypoxic or ischemic injury, we also utilized tissue from individuals with minimal change disease (MCD) as an additional reference control.

Unsupervised clustering, interaction network and functional pathway enrichment analysis was performed to investigate transcriptional regulatory profiles of AKI samples, versus reference controls (Ref and MCD) samples, then compared to corresponding clinical data. Our study will contribute to the current body of literature by adding knowledge of miRNA-mRNA associations in AKI, supplementing available KPMP data and providing insight into transcriptomic regulation of AKI which may be explored in future investigations. Further, we sought to understand the limitations of reference nephrectomy tissue as a reference comparator in acute tubular injury by comparing signatures to those of MCD samples.

## Methods

### Human subjects and histopathology

This study was approved by the Indiana University Institutional Review Board (IRB, #1601431846). Per clinical records, AKI biopsies were generally performed in cases of severe or unresolving AKI. MCD biopsies were performed for diagnostic evaluation of proteinuria (minimal change disease) or hematuria (thin basement membrane disease), but without known AKI or CKD progression. All samples were obtained in the course of routine clinical care. Kidney biopsy samples were obtained from a total of 60 individuals across three conditions: 1) MCD: individuals with the diagnosis of minimal change disease/thin basement membrane disease 2) Ref: reference and tumor nephrectomy samples and 3) AKI: individuals with a clinical diagnosis of AKI and primary histopathologic diagnosis of acute tubular necrosis (ATN) or acute interstitial nephritis (AIN). Subjects with AKI were screened from the Biopsy Biobank Cohort of Indiana (BBCI) as having a primary histopathologic diagnosis of either ATN or AIN. The clinical presence of AKI meeting a minimum of AKIN stage 1 criteria was confirmed in the electronic health record.^27^ Subjects were excluded if they had rapid reversal of AKI within 48 hours, had insufficient follow-up, a renal transplant, histopathologic or clinical evidence of glomerulonephritis, obsolescence of greater than 50% of glomeruli or IFTA >20%.^28^ The archived renal biopsy specimens were retrieved after cryostorage in O.C.T. at -80°C for up to 5 years.

### RNA extraction

From all renal biopsy samples, 2 cryosections (10 µM) including the entire cross-section of the tissue were placed directly into PicoPure RNA extraction buffer (Thermo Fisher, Waltham, MA). RNA was isolated using the PicoPure isolation kit according to manufacturer’s instructions with DNAse treatment and divided for RNAseq experiments.

### Whole transcriptome sequencing of ultra-low input RNA sequencing

Total RNA was evaluated for quantity and quality using an Agilent Bioanalyzer. Approximately 10 ng of total RNA were used for each sample’s library preparation. Ribosomal RNA was removed using RiboGone – Mammalian Kit protocol (Clontech, No. 634847). After rRNA depletion, double-stranded cDNA was synthesized and amplified using the SMARTer Universal Low Input RNA Kit protocol (Clontech, No. 634938), a product optimized for archived biosamples.^29^ The cDNA was sheared by Covaris, and the library was prepared and barcoded following the Ion Plus Fragment Library Kit protocol (Life Technologies, No. 4471252). Each resulting barcoded library was quantified and quality assessed by Agilent Bioanalyzer. Multiple libraries were pooled in equal molarity. Finally, 8 µL of 100 pM pooled libraries were loaded into lanes of a HiSeq 4000 (Illumina). At least 30M reads per library were generated. Depending on time of sample acquisition and analysis, single-end or paired-end sequencing strategies were used for AKI and Ref samples. All MCD samples underwent paired-end sequencing. As later described, only paired-end samples were used for downstream analysis unless otherwise specified.

### Micro-RNA sequencing

Prior to library preparation and sequencing, quality control of the total RNA from kidney tissue was performed using Agilent Bioanalyzer to assess RNA concentration and RIN scores. Approximately 100 ng of total RNA was used. MiRNA libraries were prepared using the QIASeq miRNA Library Kit (Qiagen). Each resulting indexed library was quantified and its quality assessed by Qubit and Agilent Bioanalyzer. The libraries were then loaded on to NextSeq 500 for 75bp single-read sequencing (Illumina, Inc), sequenced to a depth of 15-20M reads per library.

### Mapping and Expression

data is available in the Gene Expression Omnibus (GSE139061). *mRNA*: The obtained sequence data were first pre-processed to remove remaining SMART CDS prime sequences and 3 bp was trimmed off from read ends. The resulting reads were mapped to the human reference genome using STAR RNA-seq aligner (v2.5.2b).^30^ Duplicate reads were diagnosed with the Bioconductor package dupRadar.^31^ Gene-based expression levels were quantified with featureCounts (subread v.1.5.0)^32^ applying parameters “–Q 10”.

Total read counts mapping to an annotated gene were determined with edgeR.^33^ Most reads mapped to exonic regions; any intronic reads were not included in total read counts in our analysis.^34^ RNAseq samples included both single-end and paired-end sequencing methods. As later described, only paired-end samples were used for downstream analyses unless otherwise specified.

#### miRNA

Sequence reads were uploaded to the Qiagen GeneGlobe Data Analysis Center (https://www.qiagen.com/us/resources/geneglobe) for quality control, alignment and expression quantification. Reads were mapped to different database for mature, hairpin, noncoding RNA, mRNA and otherRNA using bowtie v1.2 (bowtie-bio.sourceforge.net/index.shtml). Read counts and UMI counts for each RNA category (mature, hairpin, piRNA, tRNA, rRNA, mRNA and otherRNA) were calculated for each sample using miRbase V21 for miRNA, and piRNABank for piRNA.

### Data Analyses

RStudio server (version 2023.03.0 Build 386)/R (version 4.3.1) were used with DESeq2 (version 1.40.2) in Bioconductor (version 3.17) for sample normalization each for mRNA and miRNA.^35^ Principal component analysis (PCA) plots were created using the ggplot2 (version 3.4.2) package.^36^

#### Quality control mRNA

Samples sequenced using the single-end method were excluded (n = 27 AKI). Given the variable RNA integrity of the archived clinical specimens used, count histograms of the remaining samples (n = 33) were visually evaluated on a per-sample basis and tested for fit against a negative binomial count distribution, an assumption of DESeq2. Samples that failed were excluded (n = 5 AKI, 5 MCD). Lastly, remaining samples (n = 23) were evaluated by PCA and one extreme outlier (n = 1 MCD) was removed (**Figure 1 A**).

**Figure 1:**
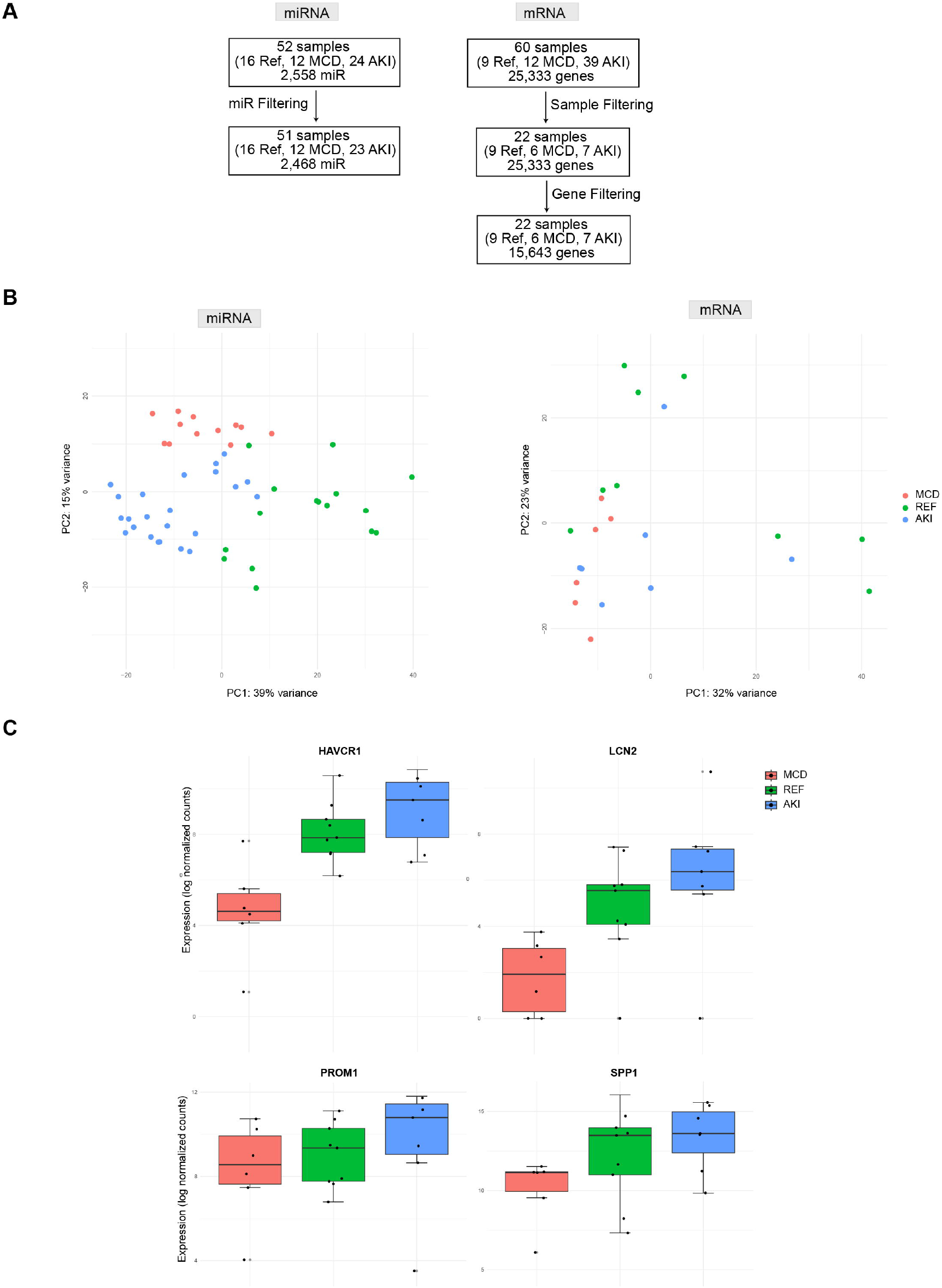
Description of dataset analyzed. A) Left panel: miRNAseq was performed on 52 kidney biopsies and after removal of 1 outlier, resulting in 51 samples used in downstream analyses (16 Ref, 12 MCD, 23 AKI) with 2,468 miRNA after data pre-processing piRNA removal (see Methods). Right panel: RNAseq was performed on 60 kidney biopsies (9 Ref, 12 MCD, 39 AKI). Sample filtering: samples sequenced using the single-end method were excluded (n = 27 AKI). Count histograms of the remaining samples (n = 33) were visually evaluated on a persample basis and tested for fit against a negative binomial count distribution, an assumption of DESeq2. Samples that failed were excluded (n = 5 AKI, 5 MCD). Lastly, remaining samples (n = 23) were evaluated by PCA and one extreme outlier (n = 1 MCD) was removed, resulting in 22 samples used in downstream analyses (9 Ref, 6 MCD, 7 AKI) and 15,643 protein-coding genes retained (see Methods). B) PCA of AKI, Ref and MCD samples each in miRNA and mRNA sequenced samples. miRNA demonstrates preserved clustering by diagnosis, with less obvious clustering in mRNA samples. C) Box-and-whisker plots of renal injury markers: *HAVCR1* (KIM1), *LCN2* (NGAL), *PROM1* and *SPP1* (Osteopontin). Note that Ref expresses injury markers similar to AKI, while MCD does not. Abbreviations: Nephrectomy (Ref), Minimal change disease (MCD), Acute kidney injury (AKI)

#### Data pre-processing

Genes (initial n = 25,333) and miRNA (n = 9,752) were excluded if they did not meet the criteria of 3 counts in at least 25% of samples in at least 1 group (n = 5,759 genes, 7,284 miRNA). Remaining non-protein coding genes defined as “Non-coding RNAs” by the HUGO Gene Nomenclature Committee (n = 3,931 genes) and piRNA (n = 238 miRNA) were removed. In total, 15,643 genes and 2,468 miRNA were retained for downstream analyses (**Figure 1 A**).

#### Identification of differentially expressed genes and microRNAs

Differentially-expressed genes and miRNAs were considered statistically significant using a likelihood ratio test and weighted Benjamini-Hochberg procedure with data-driven weights (IHW version 1.28.0) at adjusted p-value < 0.05.

#### Pathway enrichment via active subnetworks and miRNA-mRNA targetome analysis

PathfindR (version 2.1.0) using the STRING protein interaction network was used for multiple downstream pathway analyses.^37; 38^ Pheatmap (version 1.0.12) was used to generate heatmaps using Spearman dissimilarity. Pathways were considered significant at p < 0.05. Input was provided as a list of genes with log2 fold-change and p-values from DGE analysis. For miRNA pathway enrichment, miRNAs were mapped to validated gene targets using multiMiR (version 1.20.0, database version 2.3.0).^39^ MultiMiR determines validated targets using the databases miRecords, miRTarBase, and TarBase.^40-42^ For integration of miRNA and mRNA datasets, differentially-expressed miRNAs were paired with their respective differentially-expressed gene targets.

#### scRNA data analyses

The bulk mRNA seq was compared to the KPMP-HuBMAP kidney atlas (atlas).^10^ For this analysis, mRNA data was aggregated per condition, log2 transformed, and rescaled. The transformed expression of injury-associated genes was compared to a dot plot of tubular cell types in the atlas.^43^

## Results

### Dataset

Our finalized dataset included 51 samples for miRNA analyses, and 22 samples for mRNA analyses, spanning AKI (miRNA n= 23, mRNA n=7), MCD (miRNA n=12, mRNA=6), and Ref (miRNA n=16 samples, 7 of which are technical replicates from 9 subjects, mRNA n= 9). The imposition of strict quality control metrics was the reason for the different number of archived biopsy samples analyzed for mRNA and miRNA (see Methods, **Figure 1 A**). The baseline clinical and histopathologic characteristics for all subjects are provided in **Supplemental Table 1**. All subjects with AKI had a primary diagnosis of either ATN or AIN, but significant overlap in features was observed as 95% of subjects had a component of ATN and 82% had acute or chronic interstitial infiltrate on light microscopy.

### Sample clustering by miRNA and mRNA

Transcriptomic signatures of the AKI controls, Ref and MCD were compared. A principal component analysis of AKI, Ref, and MCD revealed distinct clustering of groups by miRNA expression, but less defined clustering of mRNA between the three groups (**Figure 1 B**). We assessed the genes and miRNA which contributed in the largest proportion to PC1 and PC2. Injury markers (e.g., *SPP1*), and metabolic enzyme-encoding genes (i.e, *ALDOB, CYP4A11, LRP2, CUBN*) were among the top 20 contributory genes (**Supplemental Table 2**). A similar analysis with miRNA revealed that the top contributors to clustering were: miR-19-a-5p, an important regulator of interferon-induced genes and MHC I expression in cancer, as well as epithelial-to-mesenchymal transition (EMT), ^44; 45^ and miR-1275 and miR-193-5p, both involved in the regulation of EMT (**Supplemental Table 2**).^46; 47^

### Reference nephrectomy samples demonstrate injury patterns of AKI, absent in MCD samples

Because miRNA signatures are less defined in AKI than mRNA, we first sought to characterize the mRNA signatures of our samples to describe their stage and severity of AKI. Gene expression and pathway analyses were compared between AKI, Ref and MCD. A continuum of expression was observed for the injury markers *HAVCR1* (KIM-1), *LCN2* (NGAL), *PROM1*, and *SPP1* (Osteopontin), with lowest expression seen in the MCD samples, moderate expression in Ref, and highest expression in AKI (**Figure 1 C**). Similarly, in both AKI and in Ref, we found significant upregulation of genes related to inflammation and the innate immune response (*CXCL1, CXCL8, EGR2*), hypoxia (*HMOX1*), and reparative related genes (*ATF3, EGR1*), absent in MCD (**Figure 2 A)**. We then compared DGE analysis of AKI vs. MCD with the DGE analysis of AKI vs. Ref, and Ref vs. MCD, respectively. A comparison of shared DE genes between AKI v. MCD and Ref vs MCD revealed that AKI and Ref shared 114 upregulated genes and 19 downregulated genes when each was compared to MCD (**Figure 2 B**), with several shared upregulated genes related to inflammation (*CXCL2, S100A8*), free radical generation and shared downregulated genes related to ECM reorganization (**Figure 2 C, Supplemental Tables 3**,**4**). Collectively, these analyses suggest that Ref share some transcriptomic similarities to AKI, primarily related to inflammation and innate immune responses. Given the absence of acute tubular injury patterns in MCD, comparisons of AKI to MCD afforded greater sensitivity to appreciate known pathways of AKI than comparisons of AKI to Ref.

**Figure 2:**
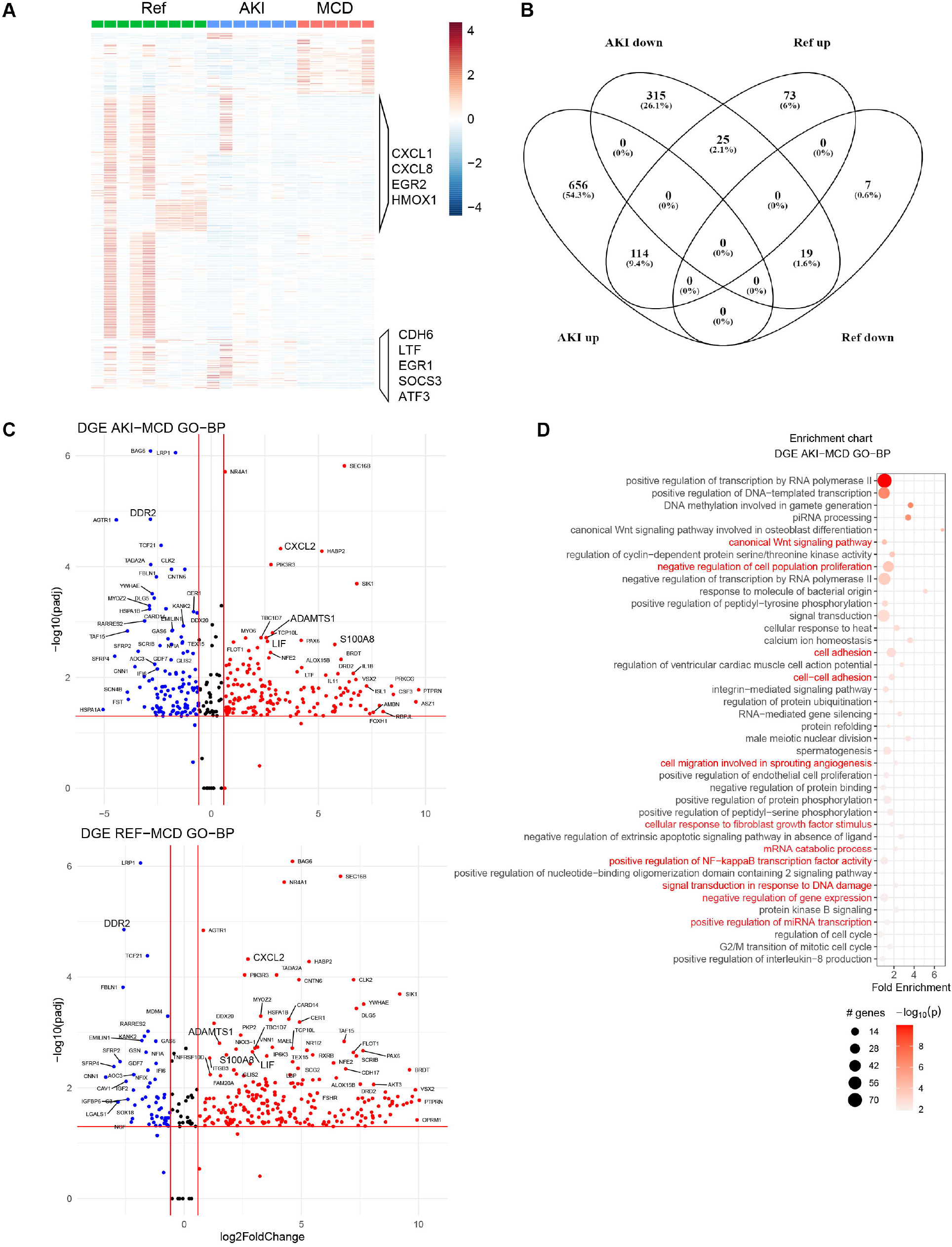
Reference nephrectomies demonstrate injury patterns, similar to AKI. A) Heatmap of gene expression patterns between Ref, AKI and MCD. AKI share similar upregulation of several injury and inflammatory genes. Values are calculated as log2FC over the mean expression across groups and scaled by row. B) Venn diagram comparing the number of shared or unique genes with a log2FC ≥1.5 and padj ≤ 0.05 for DGE of AKI vs. MCD and Ref vs. MCD. C) Volcano plots of selected DEGs enriched in the GO-BP pathway demonstrate similarities between AKI and Ref samples, selected genes in larger text reflect injury and inflammation, and are similar between AKI and Ref. Log2FC cut-offs are at -1.5 and 1.5 and – log 10 (padj <0.05). D) Pathway enrichment (GO-BP) of DEGs between AKI and MCD. Highlighted in red are examples of pathways related to EMT and fibrosis, as well as miRNA-mediated regulation of mRNA.

### The transcriptomic signature of AKI was enriched for pathways of cell adhesion and epithelial to mesenchymal transition

In a pathway enrichment analysis, we identified several interesting patterns, one of which was the modulation of genes in pathways related to adhesion and EMT (**Figure 2 D**). In our AKI cohort, biopsies were performed due to the lack of expected renal recovery in most subjects. To better understand the stage of AKI in these bulk samples, we compared their signature to the larger, cell-specific KPMP-HuBMAP kidney cell atlas.^10^ Recently, an adaptive cell state was described in cells of the proximal tubular (aPT) and thick ascending limb (aTAL), demonstrative of the equilibrium between successful and failed repair. Representative genes of this cell process include: *CDH6, PROM1, ITGB6, CCL2, CDKN1A/B* and *CDKN2A*, and transcription factors such as: *XBP1, ATF4, ATF6*, and *HIF1A* (**Figure 3 A**).^10^

**Figure 3:**
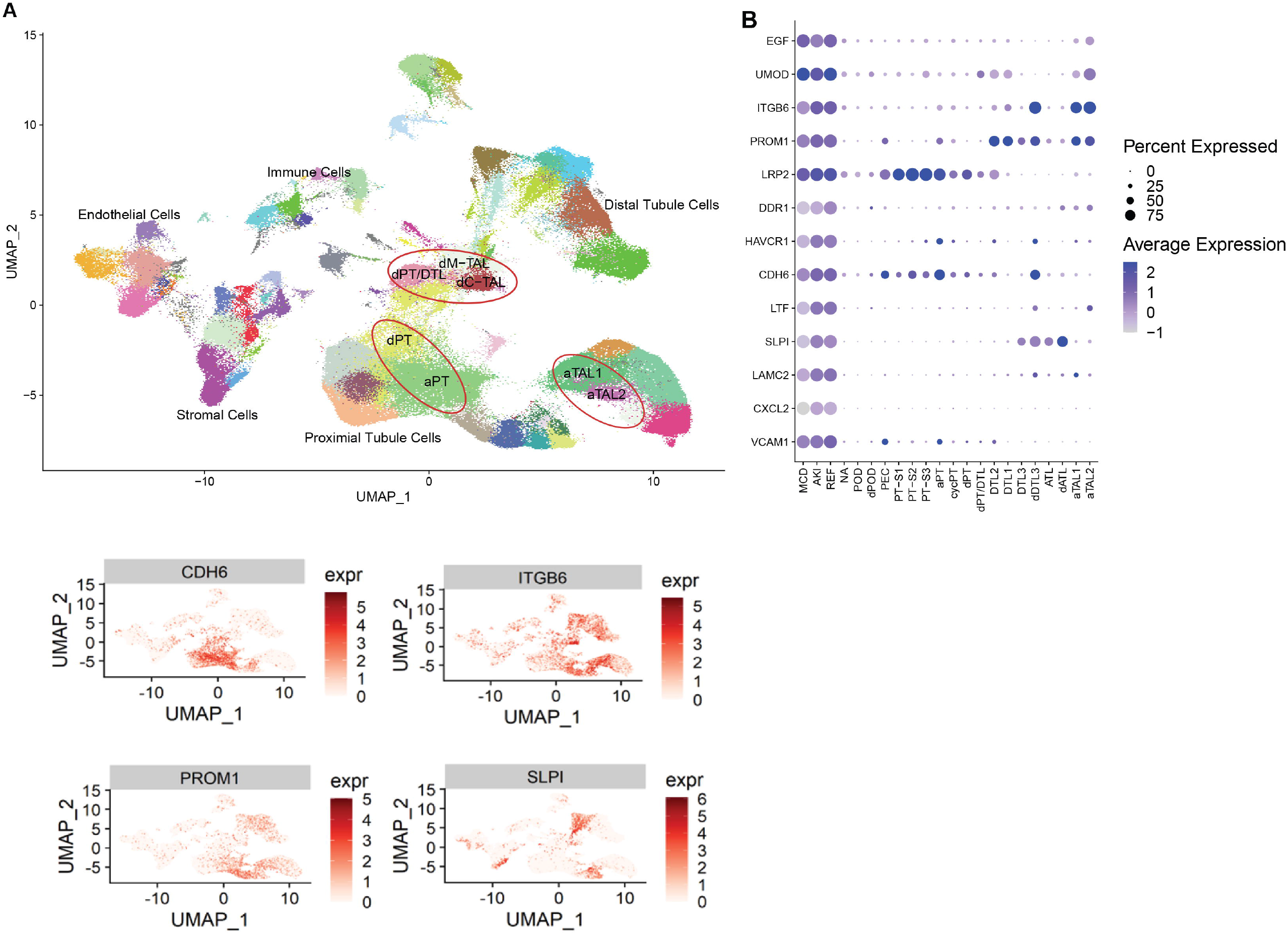
AKI samples were enriched for pathways of cell adhesion and epithelial to mesenchymal transition. A) (Upper panel) UMAP of the KPMP-HuBMAP kidney cell atlas with abbreviated labels. Circled are adaptive (aPT/aTAL) and degenerative (dPT, dTAL) cell states. In the lower panel are feature plots highlighting selected genes which characterize adaptive and degenerative cell states. B) Dot plot highlighting selected genes localized to adaptive and degenerative cell states. Note that AKI bulk data demonstrate upregulation of several these genes compared to MCD.

We identified enrichment of several marker genes of both the aPT and aTAL in our bulk AKI samples (**Figure 3 B**). For example, in AKI, *CDH6* --an aPT gene was upregulated (vs. MCD) and *ITGB6*, an upregulated gene localized to aTAL cells (vs. MCD, **Figure 3 A, B, Supplemental Table 4**). Other examples of aPT and aTAL gene expression in our data are shown in **Figure 3 B**, and other significantly upregulated genes for these cell types (padj <0.05) were identified (e.g., *S100A2, SERPINA1, CDKN1A*, see **Supplemental Table 4**). However, other marker genes such as *BMP6* or *IGF1*,^10^ were not significantly DE in our AKI samples, although we are underpowered.

### Alterations in miRNA expression

Among the pathways characteristic of AKI, we identified increased expression of miRNA and decreased expression of relevant gene targets, in addition to pathway enrichment of genes which demonstrate regulation of miRNA-mediated regulation in injured renal tissue (**Figure 2 D, Supplemental Table 5**). The top differentially expressed upregulated miRNA was miR-3170 (**Figure 4 A**) which targets *MEOX1*, a gene responsible for cell cycle regulation and progression to fibrosis.^48^ In an integrated analysis of miRNA-mRNA interactions and associated pathways, we identified groups of upregulated miRNA in our data that were associated with the downregulation of genes such as *FASLG* and *CDK7* (**Figure 4 B**), suggesting miRNA regulation of cell cycle progression and apoptotic regulation of immune cells (**Supplemental Table 5, Supplemental Table 6**). Upregulation of genes related to inflammation was also seen (e.g., *CXCL8, IL10, HMOX1, IL1B*) as shown in **Figure 4 B**. Other significant upregulated miRNA (e.g., miR-150-5p, miR-155-5p, and miR-142) are key regulators of the inflammatory immune response and fibrosis (**Figure 4 A**).^49-54^

**Figure 4:**
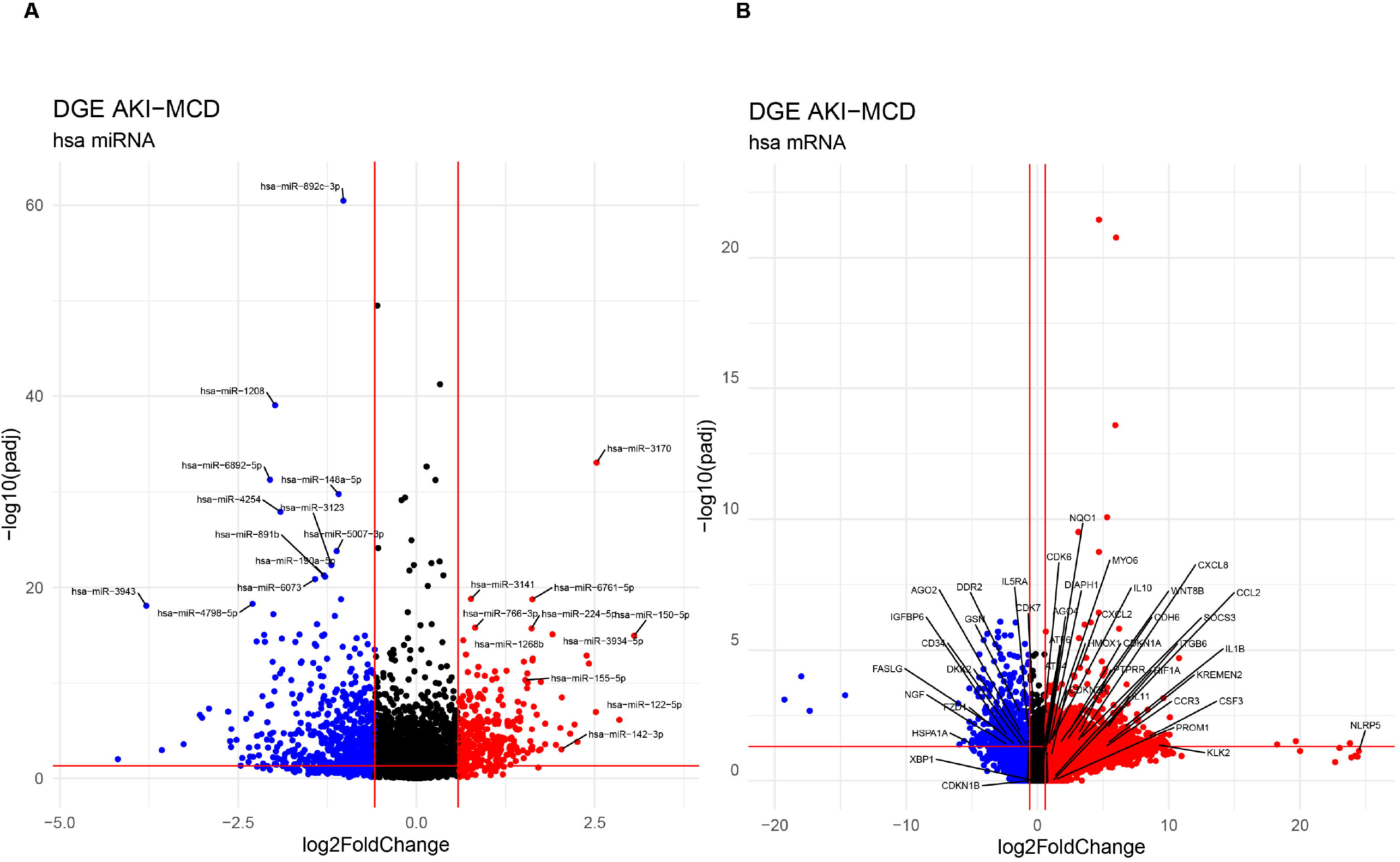
Differentially expressed miRNA and mRNA in AKI. A) Volcano plot of miRNA DGE in AKI vs. MCD. B) Volcano plot of mRNA DGE in AKI vs. MCD. Red lines denote Log2FC cut-offs are at -1.5 and 1.5 and – log 10 (padj <0.05). Red dots are upregulated and blue dots downregulated miRNA or mRNA.

We then investigated the epigenetic regulation of the genes by miRNA, associated with adaptive cell states in AKI (**Figure 5 A, B**). Notably, in AKI samples, miRNAs associated with immune recruitment and subsequent cellular and tissue remodeling, such as miR-4731 (an attenuator of EMT) and *MYO6*, and miR-146a-5p, associated with downregulation of gene targets such as *IGFBP6* (a stimulant of chemotaxis),^55; 56^ (**Figure 5 B**) were identified. Further, MiR-122-5p, predicted to bind directly to the 3’ UTR of *DDR2*, and *MMP2*, downregulated in AKI, may ultimately decrease the activity of pro-fibrotic pathways (e.g., collagenases, matrix metalloproteinases, **Figure 5 B, Supplemental Table 5, Supplemental Table 6**). Additionally, we found that miR-30a-e were significantly downregulated in AKI, while downstream targets such as *SOCS3*, were upregulated (**Supplemental Table 3**), likely inhibiting the IL-6 pathway. Other relationships were explored: *CXCL2*, upregulated in our AKI dataset is also a direct target of miR-192-5p, which was downregulated in AKI, and thus may promote inflammation. miR-150-5p, known to bind directly to *ZEB1*, may directly influence *PROM1* expression, worsening AKI. In our data, while *ZEB1* and *PROM1* were not differentially expressed in AKI vs MCD or Ref, downstream genes involved in EMT such as *ITGB6*, were upregulated in AKI (**Supplemental Table 4, Supplemental Table 6**). miR-150-5p expression was inversely correlated with *NOTCH3* expression (**Figure 5 B**). miRNA-150-5p is a target of transcription factors (e.g., *NFkB, c-Myb, b-Myb, SP1*), which upregulate miR-150 expression and exert downstream inhibition of the Notch pathway. Overall, these findings suggest complex interactions between miRs and target mRNA expression in AKI processes of inflammatory and immune responses and EMT.^57; 58^

**Figure 5:**
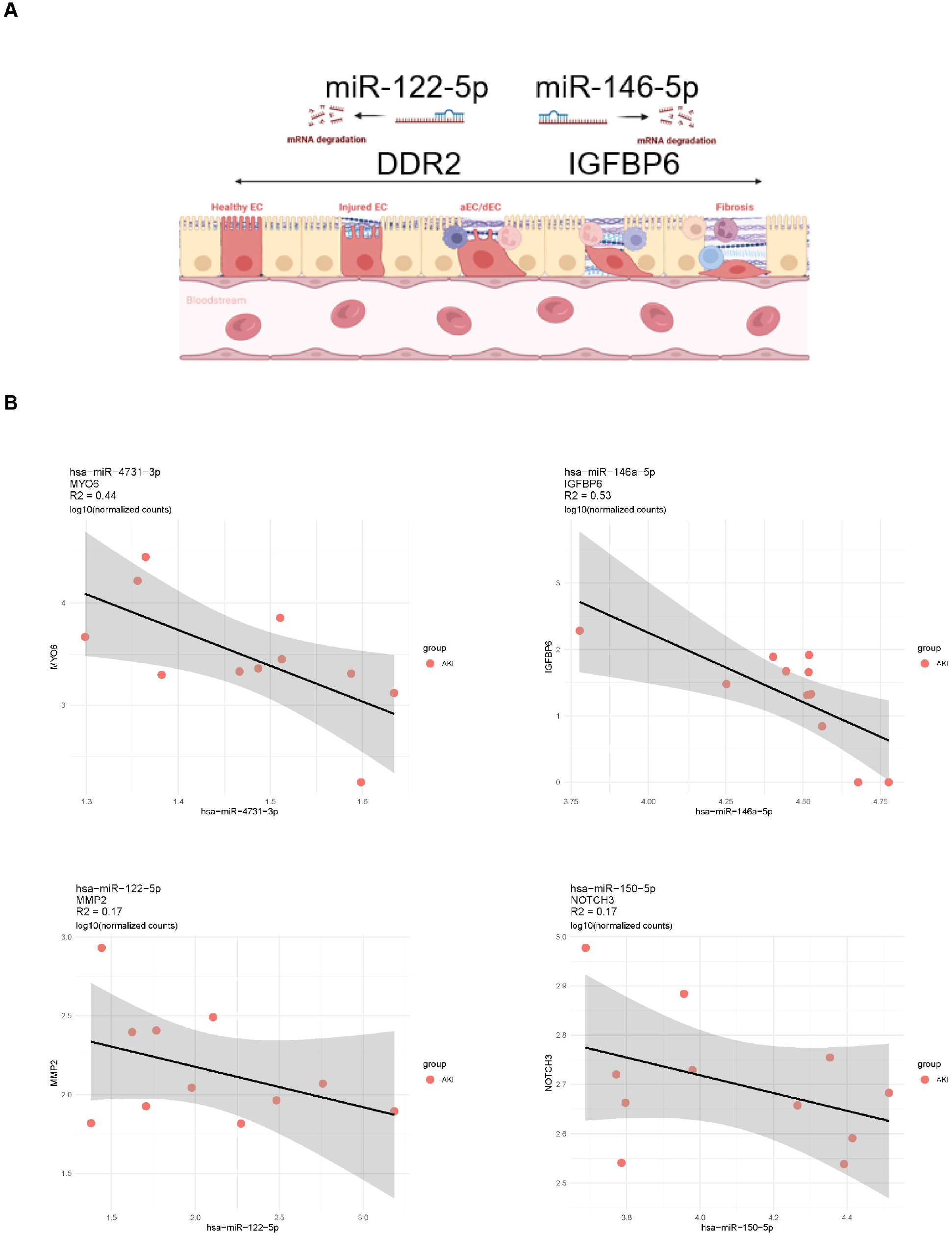
miRNA may regulate target mRNAs critical to the adaptive cell state. A) Shown in the schematic are two examples of profibrotic and anti-fibrotic miR-mRNA interactions in AKI (cells created with BioRender). B) Scatterplots of single-end AKI paired miRNA-mRNA data demonstrating inverse relationships between: miR-4731 and MYO6 and miR-146 and IGFBP6, miR-122 and MMP2, miR-150 and NOTCH3, respectively.

## Discussion

We examined the transcriptomic regulation of AKI in humans. By placing our AKI samples in the context of the KPMP-HuBMAP scRNA-seq atlas, we determined they have a strong adaptive repair molecular phenotype. We found that the genes involved in adaptive cell repair,^10^ are targeted by a network of miRNA, which regulate important pathways responsible for the inflammatory and fibrotic response in AKI, such as miR-155, a regulator of the NfkB pathway.^51^ This miRNA has been reported as a potential AKI biomarker^59^ is and also a promising therapeutic target in the treatment of AKI.^51; 60^ Similarly, we identified inverse correlations with miR-122 and miR-150 (upregulated in our AKI data), presumed to promote healing from AKI. Interestingly, similar to others, we identified a downregulation of miR-192 in AKI samples, which may lead to the initiation and propagation of tissue injury.^61^ Collectively, our findings contribute to the current body of literature and suggest roles for these miRNAs as potential therapeutic targets to mitigate failed repair trajectories in epithelial cells.^49; 50; 62; 63^

An important contribution of our work is to understand the limitations of reference nephrectomy samples as control tissue in AKI studies. Reference nephrectomies shared many alterations in immune and inflammatory pathways that were found in AKI samples. These samples, due to the nature of tissue acquisition, are prone to hypoxic injury and manifest mild injury signatures. We did not observe similar elevation of injury markers in our MCD samples. Thus, this study underscores the importance of utilizing an appropriate control when assessing tubular AKI. Although comparisons to MCD samples allowed us to uncover canonical pathways of AKI and better understand differential expression of miRNA in AKI, these samples are not appropriate to study glomerular functional tissue units. Albeit rare, living donor biopsy samples may prove an optimal control in transcriptomic studies.

Finally, we performed both mRNA and miRNA sequencing in a subset of our samples. Having a subset of matched miRNA-mRNA samples was invaluable and, we were able to detect several associations between these transcriptomes to reveal critical mediators of the AKI response. While these data contribute to several significant and novel findings when describing the epigenetic regulation of human AKI by miRNA, the analyses are primarily associative and further research in larger cohorts is needed to identify site binding targets in individuals with AKI to determine causality of these relationships.

Due to the nature of biobank tissue preservation for transcriptomics in standard O.C.T., some of the biobanked specimens had lower quality mRNA and were excluded from the analysis, leaving more miRNA samples than mRNA samples for comparison. In addition, our samples represent bulk miRNA and are not cell specific, which may also bias molecular-level expression.^64^ To help overcome both of these limitations, we compared our cohort to the larger cell-specific KPMP-HuBMAP cohort. As technology advances, cell-specific miRNA may be more readily obtained. Despite these limitations, bulk miRNA retained a remarkable ability to discriminate between AKI and the MCD and Ref groups. We therefore propose future investigations of biobanked tissue to include miRNA transcriptomics.

Overall, using computational analyses of the miRNA-mRNA targetome with our tissue-level expression data, we were able to identify relevant pathways critical to AKI pathobiology and key miRNAs important in AKI. Future studies in paired miRNA-mRNA tissue transcriptomics should employ larger datasets.

## ACKNOWLEDGMENTS

DJ was supported by Grant Number, UL1TR002529 (S. Moe and S. Wiehe, co-PIs) from the National Institutes of Health, National Center for Advancing Translational Sciences, Clinical and Translational Sciences Award, a research grant from Dialysis Clinic, Inc., and Grant 2021258 from the Doris Duke Charitable Foundation through the COVID-19 Fund to Retain Clinical Scientists collaborative grant program and was made possible through the support of Grant 62288 from the John Templeton Foundation and the IU School of Medicine Department of Medicine. The opinions expressed in this publication are those of the author(s) and do not necessarily reflect the view of the John Templeton Foundation. MTE was supported by an NIH/NCCIH award (R01AT011463-02). PCD was supported by NIH/NIDDK R01 DK107623-07.

We thank Sanjay Jain for the generous contribution of his processed single cell and single nuclear data, which helped us achieve context for many of our interesting findings.

## CONFLICT OF INTEREST

The authors have nothing to disclose.

## AUTHOR CONTRIBUTIONS

DJ, MTE, TH, PCD: Conceptualization; DJ, MTE, TD, RMF, TH, PCD: Data curation; TD, DJ, RMF, DLG: Formal analysis; JH, JL, YY: Sample curation, nephrectomy tissues; DJ, MTE, PCD, JH: Funding acquisition; All authors: Investigation; All authors: conduct of trial; KDL: Project administration; DJ, TD: Writing - original draft; All authors: Review & editing.

## DATA AVAILABILITY

Expression data is available in the Gene Expression Omnibus (GSE139061).

## Notes

https://www.ncbi.nlm.nih.gov/geo/query/acc.cgi?acc=GSE139061

### Competing Interest Statement

The authors have declared no competing interest.

